# CIDER: detecting changes in gene regulatory networks that are associated with changes in phenotype

**DOI:** 10.64898/2026.09.08.750185

**Authors:** Wooseok J. Jung, Mingyue Ding, Shu Liao, Zolboo Erdenebaatar, Michael R. Brent

## Abstract

Changes in gene regulatory networks may drive quantitative traits, or may transmit the effects of one trait, such as blood lipid level, on another, such as cardiovascular health. Yet the standard tools, differential correlation and differential network analysis, compare two discrete groups, while the contexts of interest – circulating lipids, inflammation, and blood glucose – vary continuously; applying them forces dichotomization, discarding within-trait variation. We introduce Continuous Interaction-based Differential Edge Regulation (CIDER), which tests whether a gene regulatory network edge, the relationship between a transcription factor and its target gene, varies with a continuous trait: the target gene’s expression is modeled as a function of the TF’s expression level, the trait, and their interaction, with the interaction coefficient measuring the trait dependence. To limit multiple testing, CIDER tests only the edges of a reference regulatory network. A generalized additive extension detects interactions that change the shape of the relationship, not only its slope, including forms that cannot be expressed as a difference between two correlations. In simulations it outperformed four two-group methods across sample sizes, effect sizes, and noise levels, with most of its advantage from keeping the trait continuous. In whole-blood transcriptomes from four independent human cohorts across ten quantitative health traits, CIDER identified 63 replicated cases in which a TF’s regulation of its target varies with the trait, including coupling of the glucocorticoid-receptor (NR3C1) to the granulocyte colony-stimulating-factor receptor (CSF3R) that strengthens as triglycerides rise, and a pair whose regulation reverses direction across the observed range of C-reactive protein.

## 1 INTRODUCTION

Mammalian gene expression is governed by complex regulatory networks that exhibit extraordinary plasticity. Historically, transcription factors (TFs) were conceptualized as static switches that regulate their target genes with fixed strength. However, contemporary systems biology has established that the genome operates as a highly dynamic, context-dependent system [1, 2]. The same transcription factor can strongly regulate a specific target gene in one cellular or physiological context, exert a negligible influence on the same target in a different environment, and potentially regulate it in the opposite direction (e.g., switching from an activator to a repressor) under a third set of conditions [3, 4].

In one study that compared regulatory networks across diverse human tissue and cell types, the TF-target regulatory relationships, the “edges” of the network, varied more between biological contexts than the absolute expression levels of the genes or transcription factors themselves [5]. Consequently, a TF can change which genes it regulates without accompanying change in its own expression level. Comparable TF-target regulatory rewiring accompanies disease pathogenesis, cellular differentiation, and aging [6, 7]. Genetic variation acts on the same edges: mutations in cis-regulatory regions can create, destroy, or refine a TF’s control of a target, so that a regulatory edge can differ between genotypes just as it differs between physiological states [8–10]. Regulatory relationships, therefore, must not be viewed as fixed properties of a genome, but rather as transient properties of the genome in a specific biological state. This raises the question of which regulatory relationships change together with a person’s biological state, potentially contributing to or responding to it.

The clinical and physiological variables that characterize a biological state are usually continuous. Metabolic and inflammatory markers, circulating lipids levels, body mass index, and chronological age are all inherently recorded on a continuous scale. At the population level, these biological variables tend to form a unimodal distribution rather than naturally segregating into discrete intervals or partitions. However, established tools for asking how regulatory relationships change, such as differential correlation (DC) and differential network analysis (DiNA), are fundamentally designed for the comparison of two discrete, mutually exclusive groups: cases against controls, tumor against normal tissue, or survivors against non-survivors.

Applying these established tools to a continuous trait requires dichotomizing the trait at a clinical diagnostic threshold, at the extreme tail ends of the distribution, or when no natural cutoff exists, at the sample median. The cost of this dichotomization is severe: within-group variation is entirely discarded, and granular trait signals, which often hold the key to understanding transitional disease states, are irrevocably lost [11, 12].

Two distinct bodies of work border this problem from opposite sides. DC and DiNA methods – DiffCorr, DGCA, Discordant, and DINGO – share the core biological question: how relationships between genes differ between conditions, rather than how single genes differ in isolation [13–16]. Each of these methods, however, is defined for a two-group comparison and inherits the cost of dichotomization. Conversely, gene-environment (G×E) interaction analysis in statistical genetics shares our solution: it retains continuous variables and explicitly tests them using an interaction term [17–19]. However, G×E tests whether the static effect of a genetic variant (such as a single nucleotide polymorphism) on a phenotype varies with a continuous environmental variable, not a regulatory relationship between two actively transcribed genes, which can itself change with context. Nevertheless, three lessons carry over. First, continuous variables should not be dichotomized. Second, a linear interaction alone is inadequate, as the environmental interaction can often be nonlinear. Lastly, testing interactions genome-wide incurs a heavy multiple testing burden. The established remedy is a two-step design in which candidates are screened first and interactions are only tested within that candidate set [20]. Closest to the present setting is SNiPage, which tests a SNP-by-age interaction on target gene expression, using the expression of an age-associated TF as a proxy for age [21]. In SNiPage, the variable under test is genotype and the TF stands in for the continuous context; the TF-target relationship is not itself the unit of inference, and the trait is not measured on the individual.

To our knowledge, no existing method asks whether the relationship between a TF and its target gene varies with a continuous trait while combining three properties: (a) leaving the trait continuous, (b) containing the number of tests by restricting attention to a set of candidate TF-target pairs specified in advance, and (c) testing for nonlinear interactions. Each property appears separately in prior work, as reviewed above; it is their combination that no existing method provides.

To bridge this methodological gap, we introduce CIDER (Continuous Interaction-based Differential Edge Regulation), which directly tests whether the relationship between a transcription factor and its target gene varies with a continuous trait. The target gene’s expression is regressed on the TF’s expression, the trait, and their interaction; the coefficient on the interaction term is the test statistic. Because the trait is never discretized, the quantity estimated is the TF’s regulatory effect on the target gene expression across the full range of the trait, denoted β(Trait), rather than a difference between two group estimates. CIDER takes two forms. In the linear form, β(Trait) is a straight line. CIDER-GAM replaces the product with a tensor-product interaction smooth function, allowing the relationship to change shape as a trait rises rather than only change slope. Following the two-step screening design established in G×E, candidate TF-target pairs are drawn from a reference regulatory network, so the candidate set rests on prior biological knowledge and is selected independently of the trait. We benchmark CIDER against DiffCorr, DGCA, Discordant, and DINGO in simulations varying three factors: linear vs. non-monotonic interaction form, simulated vs. real TF expression, and trait separability. We apply CIDER and CIDER-GAM to four human cohorts, LLFS, FHS, WHI, and HRS, across cardiometabolic traits, where it recovers established and previously unreported context-dependent regulation.

## 2 METHODS

### 2.1 The CIDER interaction model

CIDER tests, for each transcription factor (TF)-target gene (TG) pair, whether the regulatory relationship between the TF and its target varies with a continuous trait. We treat the trait as an effect modifier: instead of asking whether the TF and target are correlated, or whether that correlation differs between two groups, CIDER asks whether the strength or shape of the TF-TG relationship changes along the trait. The trait is the biological context in which regulation is read out.

This framing has a direct statistical consequence. Detecting context-dependent regulation reduces to testing a TF-by-trait interaction in a per-pair regression in which the target gene’s expression is the response and the TF, the trait, and their interaction are the predictors. Because the interaction is defined against the continuous trait, CIDER never discretizes it, and so avoids the loss of within-trait information incurred by the two-group methods against which we benchmark it. Each candidate TF-TG pair is tested independently, and pairs are ranked by the significance of the interaction term after multiple-testing correction.

Throughout, *X* denotes TF expression, *G* target-gene expression, and *T* the continuous trait, for a given candidate pair.

### 2.2 Linear model

In its base form, used when the interaction is assumed to act on the slope of the TF-TG relationship, CIDER fits the ordinary least squares (OLS) regression

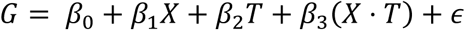

for each TF-TG pair and tests the interaction coefficient,

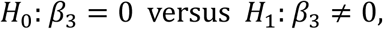

with a two-sided *t*-test on 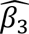. The additive terms *β*_1_*X* and *β*_2_*T* absorb any purely marginal association of the target with the TF or the trait, so that *β*_3_ isolates the interaction: a non-zero *β*_3_ means the slope of the TF-TG relationship changes with the trait, which is exactly a trait-dependent regulatory interaction. This is the model used in the linear stages (stages 1 and 2).

### 2.3 Nonlinear extension (CIDER-GAM)

The linear model captures only interactions in which the TF-TG slope changes linearly across the trait. Effect modification need not take this form: for two distinct reasons. First, at any fixed trait level the TF-TG response need not be a straight line; it may be a curve. Second, the shape of that curve, its curvature and not merely its slope, may itself change as the trait varies. To detect such interactions, CIDER generalizes the linear model to a generalized additive model (GAM), fitted with the R package mgcv [22–24], replacing each parametric term with a penalized smooth function:

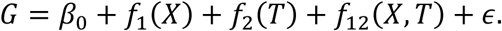

Here *f*_1_ and *f*_2_ are penalized thin plate regression splines for the main effects, the nonlinear analogues of *β*_1_*X* and *β*_2_*T*, which absorb any additive nonlinear dependence on the TF or the trait alone. The interaction is carried by *f*_12_, a tensor product interaction function, specified with mgcv’s ti() construction [25]. This construction is what makes the interaction test interpretable: because ti() excludes its marginal bases, with the corresponding marginal effects supplied separately by *f*_1_(*X*) and *f*_2_(*T*), the *f*_12_ term represents only the part of the bivariate response surface not explained by the additive main effects. It is thus the nonlinear analogue of the linear product term *X* ⋅ *T*: the pure interaction, orthogonal by construction to the main-effect smooth functions.

Each of the three smooth functions is a penalized regression spline. The two main-effect functions use low-rank thin plate spline bases, while the interaction uses a tensor product of marginal spline bases. We specify basis dimensions of *k* = 5 per term and select smoothing parameters using the restricted maximum likelihood (REML), allowing the data to determine the optimal degree of curvature. For a detailed mathematical description of the thin plate regression splines, the tensor product construction, and their respective penalty matrices, refer to the Supplementary Methods.

CIDER-GAM tests whether this interaction function is identically zero,

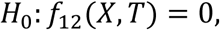

using the approximate *p*-value that mgcv reports for the ti(TF, Trait) term. These smooth-term p-values are approximate and can be mildly anticonservative under REML selection of the smoothing parameter [26]. This does not affect our benchmark, in which methods are compared by rank-based average precision rather than by the calibration of individual p-values; it is addressed separately for the cohort application, where valid inference matters. A significant interaction function indicates that the TF-TG response surface changes shape across the trait. Pairs whose GAM fails to converge are assigned a p-value of 1 and thus ranked worst, ensuring that non-convergence cannot inflate performance through survivorship bias.

Because a tensor-product interaction smooth function can represent a bilinear surface *X* ⋅ *T* as a special case, CIDER-GAM detects linear as well as nonlinear interactions effectively subsuming the linear model. We nonetheless keep the OLS model for the linear stages, as it is the more efficient and more directly interpretable estimator when the interaction is approximately linear. OLS is also the better-powered test in these scenarios: the linear model spends a single degree of freedom on the interaction, whereas the smooth function expends its power over shapes that are absent when the true interaction is linear.

### 2.4 Multiple testing and ranking

Within each simulated dataset, or replicate (Section 2.9), the 1600 interaction p-values from the linear model were adjusted for multiple testing by the Benjamini-Hochberg procedure, yielding interaction q-values and pairs were ranked by the q-value for the average-precision calculation. Because Benjamini-Hochberg is a monotone transformation of the raw p-values, this ranking is identical to ranking by the raw p-value, and the adjustment does not change the reported average precision. It is retained so that CIDER gives calibrated FDR control when used for discovery rather than benchmarking [27].

### 2.5 Simulation study

#### 2.5.1 Simulation design

We evaluated CIDER against alternative methods on simulated data in which the ground-truth set of differentially regulated TF-target gene (TF-TG) pairs is known by construction. The benchmark crosses two factors, the source of TF expression and the functional form of the TF-TG relationship, giving four stages (Table 1).

**Table 1.** CIDER simulation design.

| Stage | TF expression | TF-TG relationship | CIDER model |
| --- | --- | --- | --- |
| 1 | Simulated (independent Gaussian) | Linear interaction | CIDER-linear (OLS) |
| 2 | Real (GTEx whole blood) | Linear interaction | CIDER linear (OLS) |
| 3 | Simulated (independent Gaussian) | Nonlinear interaction (four forms) | CIDER-GAM |
| 4 | Real (GTEx whole blood) | Nonlinear interaction (four forms) | CIDER-GAM |

Stages 1 and 2 test detection of a linear TF-by-trait interaction; stages 3 and 4 test nonlinear interaction surfaces that are invisible to correlation-based methods. The simulated-TF stages (1 and 3) isolate the statistical model from the covariance structure of real expression data, while the real-TF stages (2 and 4) confirm that the conclusions hold under empirical whole blood expression. Each stage was additionally run at two trait-separability settings.

Every stage uses the same fixed dimensions: 20 TFs and 80 TGs, giving 1600 candidate TF-TG pairs, of which 10 are true positives.

#### 2.5.2 TF expression

##### Synthetic TFs (stages 1 and 3)

The expression levels of 20 TFs were drawn independently from a standard multivariate normal,

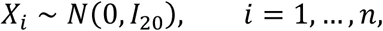

so that each TF is marginally *N*(0,1) and the TFs are mutually independent with unit variance. The quadratic interaction surface of stages 3 and 4 (Section 2.8) relies on this unit variance to simulate a pure interaction.

##### Real TFs (stages 2 and 4)

Expression levels of TFs were sampled from GTEx v10 whole blood expression (genes with ≥ 3 CPM in ≥ 25% of samples; log_10_TMM-CPM) [28]. For each replicate, the gene set was restricted to the 834 transcription factors on the Lambert list [29], 20 TFs were drawn at random without replacement, *n* donors were drawn at random without replacement from the 803 available, and each TF was standardized to zero mean and unit variance. Because sampling operates on GTEx individuals, drawing each selected donor’s entire expression vector, the natural covariance structure among TFs in a real human population is preserved: the TFs are correlated as they are *in vivo*, rather than independent as in stages 1 and 3. As the TF panel, donor set, and true-positive assignments are redrawn inside the replicate loop, variability in the choice of TF panel appears as replicate-to-replicate variation rather than being held fixed. In all four stages the trait is the simulated mixture of Section 2.6 and the target genes are generated from the models of Section 2.8; in stages 2 and 4 it is therefore only the TF expression, and with it the covariance among TFs, that is real.

### 2.6 Quantitative trait and its separability

The continuous trait was drawn from a balanced two-component Gaussian mixture,

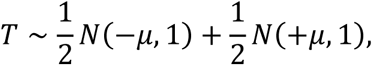

with equal mixing weights, unit component variance *σ*^2^, and symmetric component means ±*μ*. The two components represent latent subpopulations, and *μ* controls how separable they are. The trait is symmetric about zero and is bimodal if and only if *μ* > *σ* [30]. We evaluated two settings: a high-separability condition ( *μ* = 1.5) and a low-separability condition ( *μ* = 0.5). At high separability, the density is bimodal, with modes at ±1.46 and its antimode at zero; at low separability, the density is unimodal because the two components produce no trough.

The separability setting is how we probe the central methodological claim of this work. CIDER uses the trait untransformed, whereas the alternative methods cannot and must instead operate on a dichotomized trait. How well dichotomization recovers the latent structure is exactly what *μ* governs, so comparing methods across *μ* isolates the cost of discretization from any other difference between the models.

### 2.7 Trait dichotomization for the alternative methods

DiffCorr, DGCA, Discordant, and DINGO all require two discrete groups, so we split the trait at its sample median,

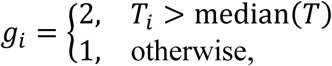

giving approximately balanced groups (*n*_1_ ≈ *n*_2_) in every replicate. The median split is not the same partition as the latent mixture-component assignment, and the two diverge most under low separability. We nonetheless regard the median split as the appropriate input for these methods: an analyst without oracle knowledge of the mixture structure has no basis for any other partition, and supplying the true component labels would credit the methods with information that is unavailable in practice. CIDER was given the continuous trait throughout and no method received the component labels.

#### 2.7.1 CIDER-discrete

To separate the cost of dichotomizing the trait from the choice of statistical model, we added a fifth method, CIDER-discrete, a variant of CIDER that fits the same model (Section 2.2) with the continuous trait *T* replaced by the binary group indicator *g* defined above. CIDER and CIDER-discrete differ only in how the trait is represented. CIDER-discrete and the four alternative methods differ only in the statistics computed from a common binary split. Comparing CIDER with CIDER-discrete therefore isolates the cost of dichotomizing the trait, and comparing CIDER-discrete with the alternative methods isolates the choice of statistical model with the trait representation held fixed.

### 2.8 Target gene expression

In every stage, 10 TGs were designated true positives and paired one-to-one with 10 randomly chosen TFs; the remaining TGs were generated as background. Each background TG carrying a TF term was paired with a TF chosen uniformly at random from the 20 TFs. Throughout, *X* denotes a TF’s expression, *T* the trait, *β* the effect size, and *ε* ∼ *N*(0, *σ*^2^) the noise. Each individual’s trait value is an independent draw from the mixture distribution of Section 2.6.

#### Linear stages 1 and 2

True-positive TGs were generated with a genuine linear TF-by-trait interaction alongside matched main effects,

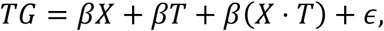

all three terms sharing the coefficient *β*. The defining signal is the multiplicative *X* ⋅ *T* term, which is what CIDER-linear tests through the OLS TF:trait coefficient. Background TGs carry no interaction; each was assigned one of four structures uniformly at random,

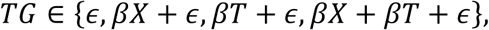

So a background gene may carry a TF main effect, a trait main effect, or both, but never an interaction. This is deliberate: a method must distinguish a change in the regulatory relationship from a mere marginal association with either variable, and a background set of pure noise would not test that.

#### Nonlinear stages 3 and 4

True-positive TGs were generated from a pure trait-dependent nonlinear interaction with no additive main-effect terms, *TG* = *β* ⋅ *f*(*X*, *T*) + *ε*. Four interaction surfaces were used, one per run, each built so that the shape of the TF-TG relationship depends on the trait (Table 2).

**Table 2.**
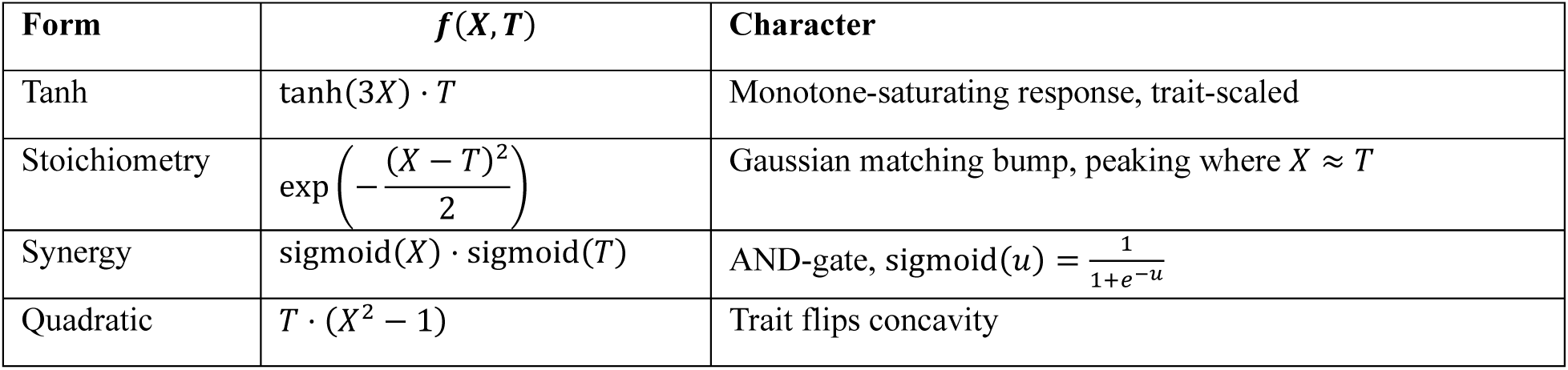
Nonlinear interaction surfaces for simulation stages 3 and 4.

| Form | $f(X, T)$ | Character |
| --- | --- | --- |
| Tanh | $\tanh(3X) \cdot T$ | Monotone-saturating response, trait-scaled |
| Stoichiometry | $\exp\left(-\frac{(X - T)^2}{2}\right)$ | Gaussian matching bump, peaking where $X \approx T$ |
| Synergy | $\text{sigmoid}(X) \cdot \text{sigmoid}(T)$ | AND-gate, $\text{sigmoid}(u) = \frac{1}{1+e^{-u}}$ |
| Quadratic | $T \cdot (X^2 - 1)$ | Trait flips concavity |

The quadratic form is the key discriminating case. With standardized TF expression, E[*X*^2^] = 1, so the −1 centering makes *T*(*X*^2^ − 1) a pure interaction with no marginal quadratic main effect, and the within-group Pearson correlation is approximately zero in both halves of the trait. Correlation-based methods are therefore blind to this form by construction, whereas a GAM can detect the concave-to-convex flip. In stage 4, real GTEx TFs are non-Gaussian, so *X*^2^ − 1 is not exactly mean-zero after standardization and a small amount of linear signal leaks; correlation-based methods consequently perform slightly above chance on this form under real TF expression.

Background TGs in the nonlinear stages were assigned to one of four structures uniformly at random,

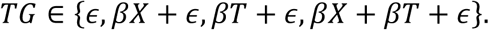

### 2.9 Simulation grid and evaluation

Each combination of stage, interaction form, and separability setting was evaluated over a grid of sample size × effect size × noise.

Sample sizes took ten values whose logarithms are evenly spaced: {10, 16, 27, 46, 77, 129, 215, 359, 599, 1000}; in the real-TF stages the largest tier was capped at 803, the number of available GTEx whole blood donors. Effect size *β* and noise *σ* were each varied over ten log-spaced values spanning [0.1,10] in the full grid. The reported benchmark uses a core subset, nine pairs of the central three values of each, *β*, *σ* ∈ {0.464, 0.774, 1.292}, crossed with all ten sample sizes, giving 90 configurations per (stage, form, separability). We call each of the nine (*β*, *σ*) combinations a design scenario and each (*β*, *σ*, *n*) combination a block. This subset spans the informative regime, where the methods are neither uniformly saturated nor uniformly at chance, while keeping the computation tractable.

Fifty replicate simulations were run per (*β*, *σ*, *n*) block, each drawing fresh TF expression, trait values, and true-positive assignments. DINGO, whose computational cost made this infeasible, was run at ten replicates per block and is excluded from the paired comparisons of Section 2.9.1. Runtimes for all methods are given in Supplementary Table S1.

Performance was summarized by the average precision (AP) of the 1600 candidate pairs against the 10-positive ground truth, averaged across replicates within a configuration to give a mean average precision (MAP). AP is rank-based and so compares methods on their ability to prioritize truly differential pairs, not on the calibration of their significance measures. Concretely, AP is computed from the ranked list: at the rank of each true positive, the precision, the fraction of pairs at or above that rank that are true positives, is recorded, and AP is the mean of these ten values. If the ten true positives occupy the top ten ranks, each contributes a precision of 1; if the tenth-ranked true positive instead sits at rank 40, its contribution is 10/40 = 0.25. AP lies between 0 and 1: a ranking that places all 10 true positives above all 1590 background pairs gives AP = 1, while ranking at random gives 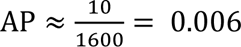. The random-ranking value follows because, under a random ordering, the expected precision at any rank equals the prevalence of true positives, 10 of 1600. Unlike the area under a receiver-operating-characteristic curve, whose null value is 0.5, the null value of AP here is close to zero, because it depends on the proportion of positives.

#### 2.9.1 Comparing methods across the design

Every method was scored on the same simulated datasets, so for a given block and replicate the average precisions of two methods form a matched pair, and the difference *d* = AP_CIDER_ − AP_alternative_ is computed on identical data. Differencing within a replicate removes the variance contributed by the dataset itself and isolates the component attributable to the method.

We tested whether these differences are positive with a Wilcoxon signed-rank test stratified by block, so that methods are compared only within a homogeneous (*β*, *σ*, *n*) group and never across blocks that differ by design. Pairs with *d* = 0 carry no information for a signed-rank test and are dropped before ranking; we call the pairs that remain *informative.* Most dropped pairs arise at the larger sample sizes, where both methods reach AP = 1.0 and the comparison saturates, so the test reflects the regime in which the methods differ. Each block contributes its own rank sum and its own null variance, the latter computed with the exact correction for ties in |*d*|. Because the blocks are independent simulations, their variances add, and the blocks combine into a single standardized statistic and a single one-sided p-value for each alternative method. One-sidedness is pre-specified by the hypothesis, that CIDER is at least as good.

At the population level, “CIDER beats the best alternative” is the same claim as “CIDER beats every alternative,” so we combined across alternatives by the intersection-union principle [31], reporting the largest of the per-alternative p-values. This needs no multiplicity correction across alternatives, avoids the selection bias of testing against whichever method happened to perform worst, and is conservative, because the single hardest competitor sets the bar. The alternative attaining the maximum is reported as the best alternative. For the Discordant method, paired replicate in which its EM algorithm failed to converge (yielding AP = 0) were dropped, so the comparison with Discordant is conditional on convergence.

Results are reported at two levels of aggregation. Within a scenario – one effect-size-noise combination (Section 2.9) – the blocks are the ten sample sizes, giving one p-value per (*β*, *σ*) scenario, which we adjust across the nine scenarios of a stage by the Bonferroni correction. Pooling all 90 blocks of a stage gives a single p-value for the design as a whole.

#### 2.9.2 Effect sizes

With 500 paired replicates behind each scenario, p-values are essentially zero wherever a real difference exists, so the effect sizes carry the quantitative message. We report three. The **peak MAP gap** is the largest value over the sample-size grid of 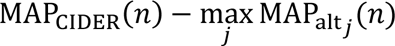, reported together with the sample size at which it occurs; because the maximum over alternatives makes the gap at each sample size the gap to the best alternative, equivalently the minimum of the pairwise gaps, the peak gap is conservative with respect to the field of competitors; taking its maximum over the ten sample sizes then reports the height of CIDER’s advantage where that advantage is largest. The **integrated gap** is the same difference integrated over log_10_ *n* by the trapezoidal rule, that is, the signed area between CIDER’s MAP curve and the upper envelope of the alternatives on a plot of MAP against log sample size. Integration is on the log scale because the grid is log-spaced and because the largest sample sizes lie in the saturated region, where both methods are at 1.0 and the gap contributes nothing; unlike the peak gap, the integrated gap is negative when CIDER trails over most of the curve. The **win rate** is the fraction of informative paired replicates in which CIDER scores higher than the best alternative. Integrated gaps are additive across nested comparisons, whereas peak gaps, being maxima attained at different sample sizes, are not; the decomposition in Section 3.3 therefore uses integrated gaps.

### 2.10 Alternative methods

We compared CIDER against four established methods for differential correlation and differential network analysis: DiffCorr, DGCA, Discordant, and DINGO [13–16]. All four are designed for comparisons between two discrete conditions and so cannot operate directly on a continuous trait. This is intrinsic to the methods, not a limitation of their implementations: each is defined by a contrast between two correlation or partial-correlation matrices, one estimated within each group, so a two-group partition is part of the estimand rather than of the interface, and there is no continuous generalization of these statistics that leaves the underlying method intact. To enable comparison, we binarized the continuous trait into low- and high-trait groups by splitting at the median. Briefly: DiffCorr and DGCA both test the difference of Fisher-transformed within-group correlations, differing in how pairs are supplied and how multiple testing is handled; Discordant fits a mixture model to paired correlation z-scores and returns a posterior probability of differential association; and DINGO decomposes each group’s partial-correlation network into shared and group-specific components, scoring edges by a standardized differential score. Complete specifications of all four methods, including implementation settings and equations, are given in Supplementary Methods.

### 2.11 Application to human cohort data

#### 2.11.1 Cohorts and expression data

We applied CIDER to whole-blood transcriptomes from four independent human population cohorts: the Framingham Heart Study (FHS), the Long Life Family Study (LLFS), the Health and Retirement Study (HRS), and the Women’s Health Initiative (WHI) [32–36]. Two of the cohorts contribute more than one sampling: FHS is represented by two examination cycles (exams 2 and 9) and LLFS by two visits (visits 1 and 2). This structure is central to the replication design (Section 2.11.5): the two FHS exams are drawn from first-degree relatives and the two LLFS visits re-sample the same individuals, so neither pairing provides genetically independent replication. After quality control, expression was available for *n* = 1971 for FHS exam 2, *n* = 715 for FHS exam 9, *n* = 1833 for LLFS visit 1, *n* = 1254 for LLFS visit 2, *n* = 3748 for HRS, and *n* = 1362 for WHI.

Gene expression was variance-stabilize transformed following NIH TOPMed pipeline freeze 10. For each candidate TF-target pair (the candidate set is defined in Section 2.11.3), the TF’s and the target’s expression values were standardized to zero mean and unit variance within the analysis stratum, one cohort exam or visit, before fitting, so that the fitted coefficients are expressed in within-stratum standard-deviation units.

#### 2.11.2 Quantitative traits

We tested ten quantitative cardiometabolic traits spanning four physiological axes: a lipid axis (total cholesterol, HDL cholesterol, LDL cholesterol, triglycerides), a glycemic axis (fasting glucose, fasting insulin, HOMA-IR), a renal axis (serum creatinine, estimated glomerular filtration rate), and C-reactive protein (CRP). Several traits are deterministic functions of others measured on the same participants and therefore do not constitute independent measurements: LDL was computed by the Friedewald estimate, LDL = total cholesterol − HDL − triglycerides/5, in mg/dL, HOMA-IR is fasting glucose × fasting insulin/405, and eGFR is a function of serum creatinine computed with the CKD-EPI creatinine equation [37–39]. Accordingly, an edge recovered for more than one trait within an axis is counted once as a single finding for that axis, while recovery across independent axes is treated as notable (Section 2.11.5).

#### 2.11.3 Reference networks and candidate edges

Rather than testing all TF-target pairs, CIDER tests only the edges of a reference regulatory network (Section 2.1), which restricts the interaction test to pairs supported by prior evidence of direct regulation and thereby limits the multiple-testing burden. We used the whole-blood METANet and whole-blood tissue-specific METANet as reference networks [40] [CITE METANet preprint]. Within each network, we took the top-500 highest-ranked candidate edges.

The two networks are analyzed as separate experiments, each with its own permutation calibration and its own Benjamini-Hochberg family, and are never pooled. Because the candidate sets are selected independently within each network, the two selections overlap only incidentally; an edge recovered in both networks is the same edge tested on the same individuals against the same trait and is counted as one finding, not two, and an edge appearing in only one network is not thereby weaker. Because the candidate set is chosen without reference to any trait, restricting it does not bias the interaction test; it is hypothesis-driven prioritization for multiplicity control, not a filter on the results.

#### 2.11.4 Interaction model and mixed-model extension for related individuals (CIDER-GAMM)

For each candidate edge and each trait we fitted the CIDER interaction model of Section 2.2 (linear form) and the CIDER-GAM model of Section 2.3 (nonlinear form), with the target gene as the response and the transcription factor, the trait, and their interaction as the terms of interest. To adjust for non-regulatory sources of covariation, the models additionally included age, age squared, sex, the top ten genetic principal components, the top ten expression principal components, red-blood-cell count, platelet count, and the white-blood-cell differential (neutrophils, lymphocytes, monocytes, eosinophils, and basophils), together with cohort-specific technical and batch covariates. The substantive covariates were applied uniformly across cohorts, so that the same adjustments enter discovery and replication, while the technical and batch terms are specified per cohort. Adjusting for the complete blood count (the red-blood-cell and platelet counts and the leukocyte differential) controls each interaction for the sample’s measured blood-cell composition directly, so that a discovered edge reflects trait-varying regulation rather than a trait-associated shift in cell-type proportions; the ten expression principal components further protect against cell-composition variation that the differential does not resolve, such a leukocyte subsets and activation states, and against residual technical variation; population structure is adjusted separately, by the ten genetic principal components.

Two cohorts contain related individuals (FHS, whose exams sample relatives; LLFS, whose design enrolls families), which violates the independence assumption of ordinary least squares. For these cohorts we used the mixed-model form of CIDER-GAM, denoted CIDER-GAMM (Generalized Additive Mixed Model), which augments the fixed-effect structure above with a random effect whose covariance is proportional to a genetic relatedness matrix derived from pedigree, fitted with mgcv. The fixed-effect structure remains identical to the unrelated-sample model, so the same interaction hypothesis is tested and only the residual covariances differ.

#### 2.11.5 Discovery and cross-cohort replication

Edges were required to replicate across genetically independent cohorts under a two-stage, round-robin procedure. In the discovery stage, each candidate edge was tested for a given trait in one cohort and retained at a per-trait Benjamini-Hochberg threshold of *q* < 0.05. In the replication stage, each discovered edge was tested in a genetically independent cohort and required to pass a Bonferroni-corrected threshold of 0.05/(*kMD*) with the same sign of the interaction coefficient as in discovery. In the Bonferroni threshold, k is the number of candidate edges (*q* < 0.05 in discovery), M the number of candidate replication cohorts, and D the number of discovery cohorts. Requiring the sign to match ensures that replication reflects the same direction of effect modification rather than unrelated significant interactions at the same edge.

The procedure is run round-robin, with each cohort serving in turn as the discovery cohort and the remaining cohorts as candidate replication cohorts. Replication is never drawn from within a cohort family: FHS exam 2 and exam 9 are first-degree relatives, and LLFS visits 1 and 2 re-sample the same participants, so neither can replicate the other; admissible replication must come from a different cohort. A (transcription factor, target, trait) triplet is reported only when it is both discovered and replicated under these rules.

## 3 RESULTS

### 3.1 CIDER estimates a trait-dependent regulatory effect and distinguishes four classes of trait dependence

Differential correlation and differential network analysis have established that the relationship between a transcription factor and its target can differ between two groups defined by a binary variable, by estimating one TF-target slope or correlation within each group (Fig. 1A, blue and red fits). Most physiological contexts of interest, however, are continuous, and applying these methods to such a trait requires dichotomizing it. CIDER instead estimates *β*(*T*), the regulatory effect of the TF on the target at trait value *T* (Fig. 1B), and tests whether *β*(*T*) varies with the trait.

**Figure 1.**
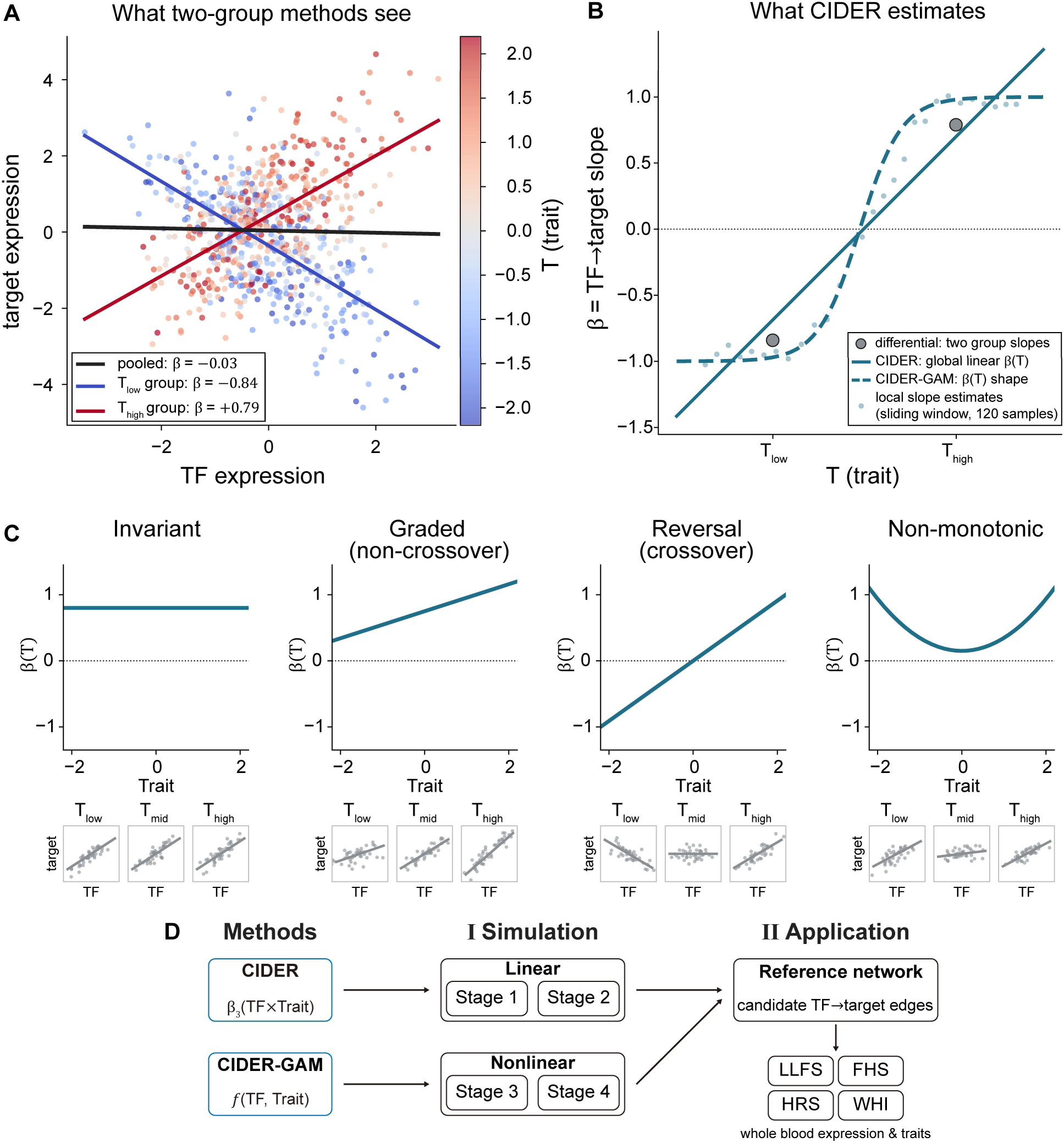
CIDER reframes differential regulation as a continuous interaction test. (A) What two-group methods see. Simulated data (n=800, SD=1.0) for one TF-target pair, colored by a continuous trait *T*. Differential correlation and differential network analysis methods require discrete groups, so *T* is split at its median and one slope is fitted per group: the low-trait (blue) and high-trait (red) slopes have opposite signs, while the overall fit (black) is flat. For these methods, detecting the difference requires dichotomizing the trait; CIDER detects it with the trait left continuous (panel B). (B) What CIDER estimates. The same data recast as *β*, the TF→target slope, against *T*. Two-group methods return two values (gray points). CIDER estimates *β*(*T*) = *β*_1_ + *β*_3_*T* over the observed range (solid line) and tests the TF×trait coefficient *β*_3_. CIDER-GAM replaces the linear interaction with a tensor-product interaction smooth function and recovers shapes the linear form cannot represent (dashed line). Light points are model-free local slopes fitted in a sliding window of 120 samples ordered on *T*, an empirical reference. (C) Typology of interactions, defined by *β*(*T*). Invariant: constant and non-zero, a real edge whose strength does not vary with the trait (the interaction null). Graded (non-crossover): monotone, single-signed, so strength changes but direction is preserved. Reversal (crossover): monotone through zero, so direction reverses across the trait range. Non-monotonic: *β*(*T*) turns around (a U-shape is shown). The first three are representable by a linear interaction, differing in whether *β*_3_ = 0 and whether *β*(*T*) crosses zero; only non-monotonic requires CIDER-GAM. Insets show the TF-target scatter and linear fit at low, mid, and high trait. (D) Study design. CIDER and CIDER-GAM are benchmarked against existing differential methods on linear (stages 1-2) and nonlinear (stages 3-4) simulations, then applied to whole-blood expression and quantitative traits in four cohorts (LLFS, FHS, HRS, WHI), with a reference network defining the candidate TF→target edges tested.

Framing the problem this way makes the possible forms of trait dependence explicit. *β*(*T*) falls into one of four classes (Fig. 1C). An invariant *β*(*T*) does not change with the trait; this is the interaction null, in which the TF-target relationship is real and non-zero and only its dependence on the trait is absent (Fig. 1C, flat line above zero). A graded *β*(*T*) changes in magnitude but keeps its sign, so the relationship strengthens or weakens without changing direction (Fig. 1C, increasing line that does not cross 0). A reversal, or crossover, passes through zero within the observed trait range, so the direction of the TF-target association inverts across the trait (Fig. 1C, increasing line that crosses 0). A non-monotonic *β*(*T*) turns around, whether U-shaped as in Fig. 1C, humped, or cubic-like. The first three classes are all straight lines in *T* and are representable by a linear interaction term; only the fourth is not, and it is the reason CIDER has a nonlinear form. The distinction between graded and crossover dependence is adapted from the reaction-norm and gene-environment interaction literature [41, 42]; the non-monotonic class has no counterpart in a two-genotype reaction-norm.

The classes are defined by the shape of *β*(*T*), not by its functional form. Fig 1C draws the first three as straight lines because those are their simplest instance, but nothing in the definitions requires linearity: a graded *β*(*T*) may saturate or accelerate and remains graded so long as it keeps one sign, and a crossover may approach and leave zero at different rates. What separates the first three classes from the fourth is monotonicity, not linearity.

This has a direct consequence for the two forms of CIDER. The linear model represents *β*(*T*) by a straight line and therefore estimates its best linear approximation. That is sufficient to detect any monotone *β*(*T*), curved or not, because the best linear approximation to a monotone function has a non-zero slope. However, representability is not the same as power: a strongly curved monotone trait dependence can reach significance under the smooth-function test while falling short under the linear one, so the two forms can disagree in practice even within the monotone classes. What the linear model cannot do is recover the shape of the curve: a curved graded or crossover *β*(*T*) will be reported with the wrong rate of change and, for a crossover, a displaced crossing point. A non-monotonic *β*(*T*) fails differently and more severely. When the turn is symmetric about the center of the trait distribution, its best linear approximation has slope near zero, so the linear interaction test is blind to it by construction rather than merely imprecise. Detecting that class is what CIDER-GAM is for; recovering the shape of a curved monotone *β*(*T*) is a second thing it buys.

We evaluated CIDER on simulated data in which the differentially regulated pairs are known by construction, across a 2 × 2 design that crosses the functional form of the interaction with the source of TF expression, and then applied it to quantitative health traits in four cohort studies (Fig. 1D).

### 3.2 CIDER outperforms alternative methods when the interaction is linear

We benchmarked CIDER against DiffCorr, DGCA, Discordant, and DINGO on simulated data with 10 truly differentially regulated pairs among 1600 candidate TF-target pairs, so that a random ranking attains an average precision of about 0.006 (Section 2.9). This section reports the two linear stages: stage 1, in which TFs’ expression levels are drawn from independent Gaussians, and stage 2, in which entire gene expression profiles of whole blood are sampled from GTEx donors. The four alternative methods received the trait split at its median, the partition available to an analyst without knowledge of the underlying mixture (Section 2.7).

In stage 1 under high trait separability, CIDER attained a higher mean average precision (MAP) than every alternative across sample sizes, effect sizes, and noise levels (Fig. 2A, B). Testing CIDER against the best alternative in each design scenario with the stratified paired Wilcoxon signed-rank test (Section 2.9.1), the advantage was significant in all nine scenarios after Bonferroni correction, with a weakest adjusted p-value of 2.7 × 10^-24^; the best alternative was DGCA in eight scenarios and DiffCorr in one. The peak MAP gap ranged from 0.23 to 0.75 across scenarios (median 0.51), and CIDER scored higher than the best alternative in at least 92% of informative paired replicates in every scenario (Table 3). The largest gaps and the smallest p-values occur in different scenarios, for a mechanical reason rather than a substantive one: where the effect is strong and the noise low, both methods saturate at AP = 1.0, so most paired replicates tie and are dropped, and with few informative pairs the signed-rank evidence weakens even though CIDER wins essentially all of the pairs that remain (win rates 0.92-1.00; Table 3).

**Figure 2.**
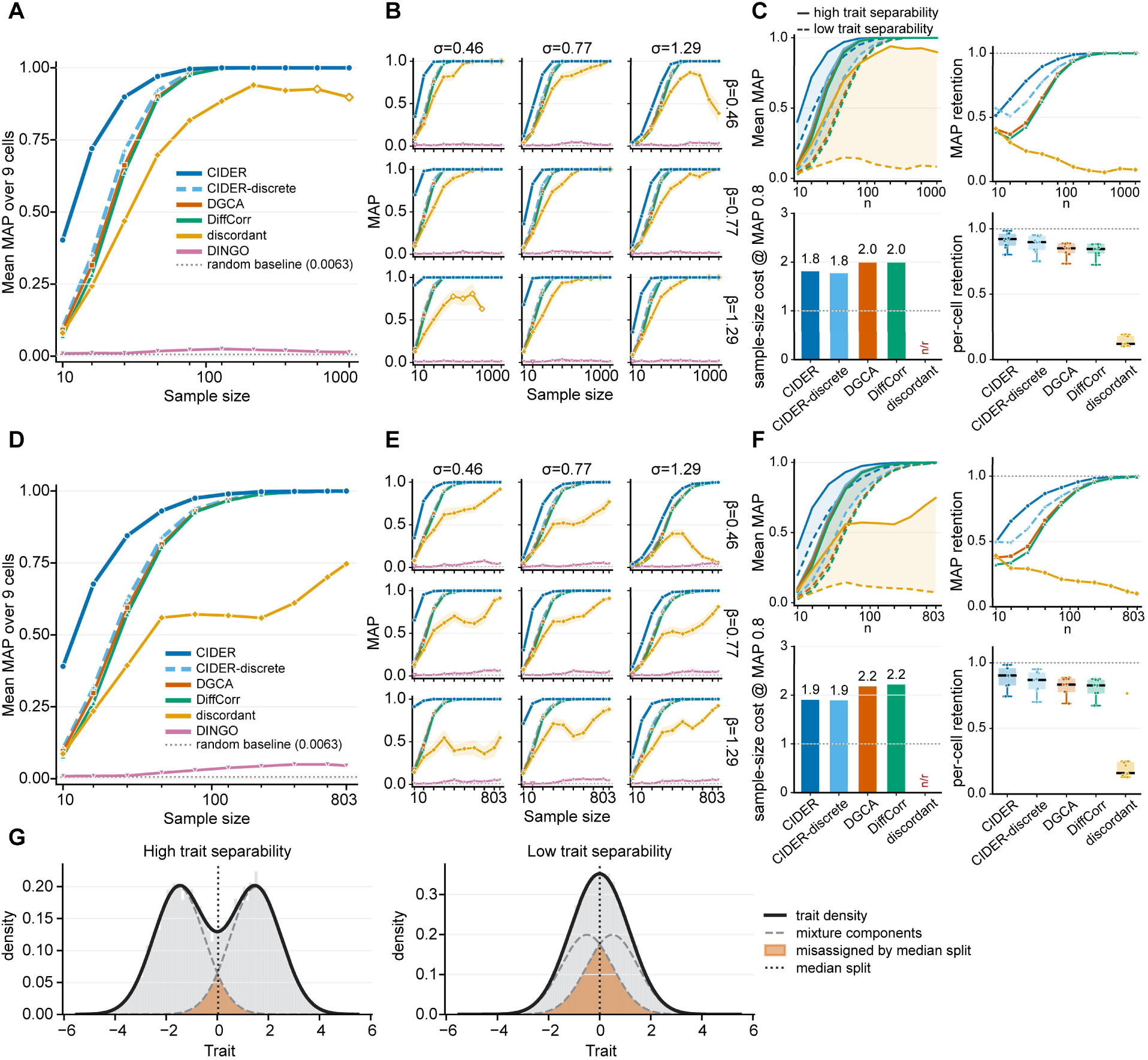
Evaluation on the linear simulations. Performance of CIDER, CIDER-discrete, DiffCorr, DGCA, Discordant, and DINGO on simulation stages 1 and 2, which have linear ground-truth interactions. (**A**) Overall performance in stage 1, the mean of the MAP scores across nine simulation scenarios. (**B**) Performance in each of the nine simulation scenarios. (**C**) Evaluation metrics comparing performance under high versus low trait separability. **Top-left**: mean MAP plotted against sample size under high trait separability (solid line) and low trait separability (dashed line). Shaded band marks the separability penalty. **Top-right**: performance retention as a ratio of MAP at low-trait separability to MAP at high-trait separability. **Bottom-left**: for each method, the sample size needed to reach MAP 0.8 in high- and low-trait separability settings is estimated by linear interpolation in log-n. The bars show the multiplicative extra sample size low separability requires to reach the same 0.8 MAP. “n/r” marks a method that never reaches the 0.8 target in either condition. **Bottom-right**: for each of the 9 simulation scenarios (cells), the area under the cell’s MAP-vs-log-n curve is computed for high- and low-trait separability settings and the low-versus-high retention ratio is computed. Boxplots show the distribution of these 9 per-cell ratios per method which preserves cell-to-cell variability. (**D-F**) The same evaluation for stage 2. (**G**) Distribution of the trait under high separability setting (left) and low trait separability (right); the mixture-component means are ±1.5 and ±0.5 respectively, with unit component variances in both. Under high separability, the trait is clearly bimodal; under low separability, it is unimodal, more closely resembling typical trait distributions.

**Table 3.** CIDER versus the best competing method across the stage-1 linear design under high trait separability. For each simulation scenario (3 effect sizes β × 3 noise levels σ), CIDER was compared against each alternative (DGCA, DiffCorr, Discordant) by a paired Wilcoxon signed-rank test stratified by sample size, on 50 replicates. All methods are scored on identical simulated datasets, so comparisons are paired. *p*_bonf_ is the intersection-union p-value, the maximum across the three alternatives, so each row is a worst-case comparison against the alternative hardest to beat (Best alt); Bonferroni correction is applied across the nine scenarios. Max MAP gap is the largest observed difference between CIDER and the best alternative method across sample sizes. Integrated gap is the area between CIDER’s MAP curve and the best alternative’s MAP curve. Win rate is the fraction of informative paired replicates (those with a non-zero AP difference) on which CIDER scores higher.

| $\beta$ | $\sigma$ | $p_{bonf}$ | Best alt | Max MAP gap | Integrated gap | Win rate |
| --- | --- | --- | --- | --- | --- | --- |
| 0.46 | 0.46 | 2.05E-28 | DGCA | 0.51 | 0.19 | 0.98 |
| 0.46 | 0.77 | 7.48E-35 | DGCA | 0.33 | 0.18 | 0.98 |
| 0.46 | 1.29 | 4.40E-38 | DGCA | 0.23 | 0.15 | 0.92 |
| 0.77 | 0.46 | 5.28E-26 | DGCA | 0.58 | 0.22 | 1.00 |
| 0.77 | 0.77 | 1.62E-31 | DiffCorr | 0.53 | 0.21 | 0.99 |
| 0.77 | 1.29 | 9.76E-33 | DGCA | 0.36 | 0.18 | 0.97 |
| 1.29 | 0.46 | 2.72E-24 | DiffCorr | 0.75 | 0.20 | 1.00 |
| 1.29 | 0.77 | 7.01E-27 | DiffCorr | 0.54 | 0.22 | 1.00 |
| 1.29 | 1.29 | 2.92E-29 | DGCA | 0.49 | 0.20 | 0.99 |

DiffCorr and DGCA performed almost identically throughout, since both reduce to the same Fisher z-test of a difference in correlation and differ only in how pairs are supplied and how multiple testing is handled (Supplementary Methods); the equivalence is documented in Supplementary Fig. S1.

### 3.3 Most of CIDER’s advantage comes from keeping the trait continuous

CIDER differs from the alternative methods in two ways: it models the target gene’s expression by regression rather than contrasting within-group correlations, and it uses the continuous trait as measured rather than split into two discrete categories. To separate these, we ran CIDER-discrete, which applies the CIDER model to the same binarized trait supplied to the alternatives (Section 2.7.1). Comparing CIDER with CIDER-discrete isolates the cost of dichotomization, since nothing else differs between them; comparing CIDER-discrete with the best alternative isolates what remains once both operate on the same binary split.

Both terms favor CIDER, but by different margins. CIDER outperformed CIDER-discrete in all nine scenarios of stage 1 under high trait separability, with a weakest adjusted p-value of 1.9 × 10^-23^, peak MAP gaps of 0.21 to 0.74 (median 0.46), and win rates of 0.91 to 1.00 (Table 4). CIDER-discrete also outperformed the best alternative in all nine scenarios, but much less: peak gaps of 0.03 to 0.08 (median 0.05), win rates of 0.62 to 0.75, and a weakest adjusted p-value of 0.012 (Table 5). Both effects are therefore real, though they differ in size.

**Table 4.** CIDER versus CIDER-discrete across the stage-1 linear design under high trait separability. . For each simulation scenario (three effect sizes β × three noise levels σ), CIDER was compared against CIDER-discrete by a paired Wilcoxon signed-rank test stratified by sample size, on 50 replicates. MAP gap is the largest observed difference between CIDER and the best alternative method across sample sizes. Integrated gap is the area between the two MAP curves; win rate is the fraction of informative paired replicates on which CIDER scores higher. *p*_bonf_ is Bonferroni-corrected across the nine scenarios.

| $\beta$ | $\sigma$ | $p_{\text{bonf}}$ | Max MAP gap | Integrated gap | Win rate |
| --- | --- | --- | --- | --- | --- |
| 0.46 | 0.46 | 6.21E-27 | 0.46 | 0.17 | 0.98 |
| 0.46 | 0.77 | 1.92E-32 | 0.27 | 0.15 | 0.97 |
| 0.46 | 1.29 | 2.79E-34 | 0.21 | 0.14 | 0.91 |
| 0.77 | 0.46 | 3.68E-25 | 0.56 | 0.19 | 1.00 |
| 0.77 | 0.77 | 1.22E-28 | 0.46 | 0.17 | 0.99 |
| 0.77 | 1.29 | 2.71E-33 | 0.30 | 0.15 | 0.97 |
| 1.29 | 0.46 | 1.92E-23 | 0.74 | 0.19 | 1.00 |
| 1.29 | 0.77 | 2.34E-25 | 0.53 | 0.19 | 1.00 |
| 1.29 | 1.29 | 1.09E-26 | 0.44 | 0.16 | 0.99 |

**Table 5.** CIDER-discrete versus the best competing method across the stage-1 linear design under high trait separability. . For each scenario (three effect sizes β × three noise levels σ), CIDER-discrete was compared against each alternative (DGCA, DiffCorr, Discordant) by a paired Wilcoxon signed-rank test stratified by sample size, on 50 replicates. *p*_bonf_ is the intersection-union p-value, the maximum across the three alternatives (Best alt); Bonferroni correction is applied across the nine scenarios. MAP gap is the largest observed difference between CIDER and the best alternative method across sample sizes. Integrated gap is the area between the CIDER-discrete’s MAP curve and the best alternative’s MAP curve. Win rate is the fraction of informative paired replicates CIDER-discrete scores higher.

| $\beta$ | $\sigma$ | $p_{bonf}$ | Best alt | Max MAP gap | Integrated gap | Win rate |
| --- | --- | --- | --- | --- | --- | --- |
| 0.46 | 0.46 | 4.27E-04 | DGCA | 0.04 | 0.02 | 0.67 |
| 0.46 | 0.77 | 7.36E-09 | DGCA | 0.06 | 0.03 | 0.67 |
| 0.46 | 1.29 | 4.39E-09 | DGCA | 0.04 | 0.02 | 0.68 |
| 0.77 | 0.46 | 1.37E-04 | DGCA | 0.05 | 0.02 | 0.69 |
| 0.77 | 0.77 | 4.61E-12 | DGCA | 0.08 | 0.04 | 0.75 |
| 0.77 | 1.29 | 1.21E-06 | DGCA | 0.06 | 0.03 | 0.66 |
| 1.29 | 0.46 | 1.17E-02 | DGCA | 0.03 | 0.02 | 0.62 |
| 1.29 | 0.77 | 2.95E-03 | DGCA | 0.05 | 0.02 | 0.64 |
| 1.29 | 1.29 | 1.72E-08 | DGCA | 0.08 | 0.03 | 0.71 |

Because the integrated gap is additive across the two comparisons (Section 2.9.2), the total advantage can be apportioned between them. Under high trait separability, dichotomization accounts for a median of 89% of CIDER’s advantage over the hardest alternative across the nine scenarios of stage 1 (range 81-92%), and the choice of model accounts for the remaining 11%.

### 3.4 CIDER’s advantage grows as the trait becomes less separable

Dichotomization is least damaging when the trait has two well-separated groups to recover. We therefore repeated the benchmark with a less separable trait. The trait is a balanced two-component Gaussian mixture with unit variances and means ±*μ*, whose density is bimodal if and only if *μ* > 1 (Section 2.6). At the high-separability setting, *μ* = 1.5, the density is bimodal with modes at ±1.46 and its antimode at the median, so the median split coincides with the trough of the density (Fig. 2G, left). At the low-separability setting, *μ* = 0.5, the density is unimodal (Fig. 2G, right): the two components are still present, but there is no trough.

All methods lost performance under the less separable trait, but CIDER lost the least (Fig. 2C). Expressed as a sample-size cost, reaching a MAP of 0.8 under low separability required 1.8 times the sample size needed under high separability, the smallest penalty of any method tested. The ratio is undefined for any method that never reaches MAP 0.8 anywhere in the sample-size grid, which affects Discordant and DINGO; for those methods the cost if reported as exceeding the largest sample size tested (censored at n = 1000) rather than being omitted (Fig. 2C).

The decomposition of Section 3.3 shifts accordingly, and not in the way a simple account of dichotomization would predict. The dichotomization effect is nearly unchanged between separability settings (median integrated gap 0.172 to 0.191 in stage 1), whereas the model-form term grows roughly fourfold (0.024 to 0.106), moving the split from 89/11 to 65/35. When the median split is uninformative, in other words, the residual advantage of modeling the target gene by regression grows: a regression that estimates the target’s conditional mean and adjusts for both main effects degrades more gracefully than a contrast of two within-group correlations, whatever the trait representation.

In stage 2, where TF expression is resampled from GTEx donors, the pattern is unchanged (Fig. 2D-F). Resampling introduces correlations among TFs’ expression levels and lowers every method’s performance relative to the synthetic stage 1, but CIDER remained significantly better than the best alternative in all nine scenarios at both separability settings, with median peak gaps of 0.43 and 0.47 respectively, and the decomposition shifted from 91/9 to 75/25. DINGO performed near chance at every configuration in both linear stages and was run at a reduced replicate count because of its computational cost (Section 2.9); it is excluded from the paired comparisons and reported separately in Supplementary Table S2.

### 3.5 CIDER-GAM recovers an interaction that no contrast of correlations can represent

The linear stages ask which method best detects an interaction that all of them can, in principle, represent. Stages 3 and 4 ask what happens when the interaction takes a shape that a difference in correlation cannot express. We simulated four nonlinear interaction surfaces (hyperbolic tangent, stoichiometry, synergy, and quadratic; Fig. 3A, Section 2.8) – and analyzed them with CIDER-GAM. Main-text results are from stage 4, with real GTEx TF expression (Fig. 3B, C); stage 3 is given in Supplementary Fig. S2.

**Figure 3.**
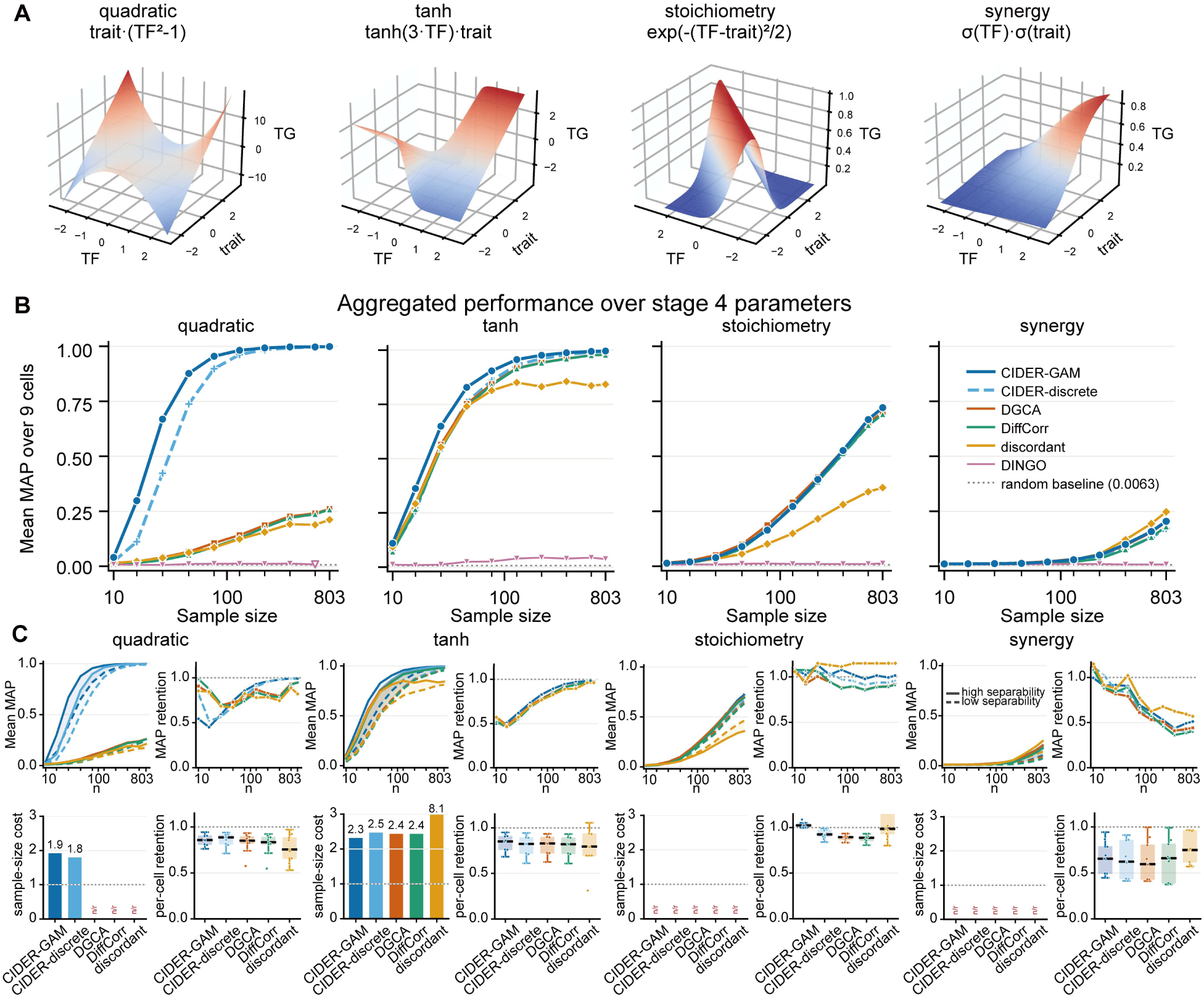
Evaluation on the nonlinear simulations. Performance of CIDER-GAM, CIDER-discrete, DiffCorr, DGCA, Discordant, and DINGO on simulation stage 4, which has four nonlinear ground-truth interaction forms. (A) The four forms, shown as target gene expression surfaces over TF expression and trait. (B) Overall performance in stage 4, the mean of the MAP scores across the nine scenarios. (C) For each form, evaluation metrics comparing performance under high versus low trait separability.

The four forms differ in a way that predicts the results, and the relevant property is not the shape of the surface itself, but what a within-group correlation can see of it. For each form we computed the Pearson correlation between TF and target expression separately below and above the trait median, the same split the correlation-based methods receive (Section 2.7), from the noise-free surface under stage-3 generative distributions (*TF* ∼ *N*(0,1); trait is generated by two-component Gaussian mixture; 2 × 10^6^ draws). Because Pearson correlation is scale-free these values depend only on the form and not on the effect size (Supplementary Table S3). For the hyperbolic-tangent surface the correlation reverses across the split, from −0.77 to +0.77, and for the stoichiometry surface from −0.60 to +0.60; in both cases the interaction shows up precisely as a change in linear association, which is the quantity these methods estimate. For the synergy surface, the correlation stays positive in both halves and does not reverse, rising from +0.53 to +0.92, because multiplying the TF response by a positive trait-dependent scale factor preserves the direction of association. For the quadratic surface the correlation is zero in expectation in both halves: TF is symmetric about zero and independent of the trait, so Cov(TF, trait ⋅ (TF^2^ − 1)) = E[trait] ⋅ E[TF^2^ − TF] = 0 within any subset of the trait. Our estimates, +0.0009 and −0.0033, depart from zero only by Monte Carlo error (SE ≈ 0.002). The TF-target relationship nonetheless inverts from concave to convex across the trait, so this form is invisible to a correlation contrast by construction.

The benchmark follows this ordering. On the quadratic form, CIDER-GAM outperformed the best alternative in all nine scenarios at both separability settings, with peak MAP gaps of 0.84 to 0.94 (median 0.89) and win rates of 0.91 to 0.99, while the alternatives remained near the 0.006 chance baseline (Fig. 3B, right). This is the largest margin anywhere in the benchmark. On the hyperbolic-tangent form, CIDER-GAM again outperformed the best alternative in every scenario, but the margin was modest. Because a tanh interaction is visible to a correlation contrast as a sign change between the trait halves, every method eventually detects it, and CIDER-GAM’s edge is concentrated at the smaller sample sizes: peak gaps of 0.08 to 0.22 (median 0.10) and win rates of 0.68 to 0.90 (Fig. 3C).

On the remaining two forms, CIDER-GAM held no advantage. On the synergy form, the methods were indistinguishable: peak gaps did not exceed 0.08, win rates ranged from 0.40 to 0.56, and no scenario reached significance at either separability setting. On the stoichiometry form, the outcome varied by scenario, with win rates from 0.26 to 0.79 and CIDER-GAM behind the best alternative in 10 of 18 scenario-by-separability combinations; because the sign of the difference is not consistent across scenarios, we report the per-scenario results and claim no advantage for either method on this form (Supplementary Table S4).

One further result separates the two axes of the benchmark. CIDER-discrete, which sees only the median-split binary trait, also recovers the quadratic form (Fig. 3B, dashed line), and does so with a large margin over the correlation-based methods that receive the same binary trait. In the linear stages, trait representation accounts for 89% of CIDER’s advantage; on the quadratic form, model representability accounts for 88-92% of it.

### 3.6 CIDER identifies context-dependent regulation across quantitative health traits

We applied CIDER to whole-blood transcriptomes from four population cohorts (FHS, LLFS, HRS, and WHI) across ten quantitative health traits, testing candidate TF-target edges drawn from two METANet reference networks. The whole-blood METANet and the whole-blood tissue-specific METANet are analyzed as separate experiments and are never pooled. A TF-target-trait triplet is reported only if it was discovered in one cohort at per-trait Benjamini-Hochberg q < 0.05 and then cleared the 0.05/(kMD) Bonferroni threshold in a genetically independent cohort with the same sign of *β*_3_; replication between FHS exams, or between LLFS visits, is inadmissible because those strata share individuals or first-degree relatives. 36 triplets met this requirement under CIDER and 40 under CIDER-GAM, spanning 63 distinct (TF, target, trait) triplets. Every statistic reported below was fixed before any literature was consulted; the supporting biology annotates the ranked candidates and does not select or reorder them. We describe two TF-target edges in detail.

#### NR3C1→CSF3R: glucocorticoid-receptor coupling to the G-CSF receptor varies significantly with circulating triglycerides

In the whole-blood METANet, CIDER identified a regulatory relationship between NR3C1, which encodes the glucocorticoid receptor (GR), and CSF3R, which encodes the granulocyte colony-stimulating factor receptor (G-CSFR), that varies significantly with triglycerides. In LLFS visit 1, the interaction estimate was *β*_3_ = +0.031 per SD of log-triglycerides (*q* = 5.6 × 10^-3^), replicating in HRS at *β*_3_ = +0.037 (*p* = 4.7 × 10^-9^ after Bonferroni correction) (Fig. 4A). At the cohort mean, NR3C1 positively regulates CSF3R, but this coupling strengthens substantially in individuals with higher triglyceride levels (Fig. 4B).

**Figure 4.**
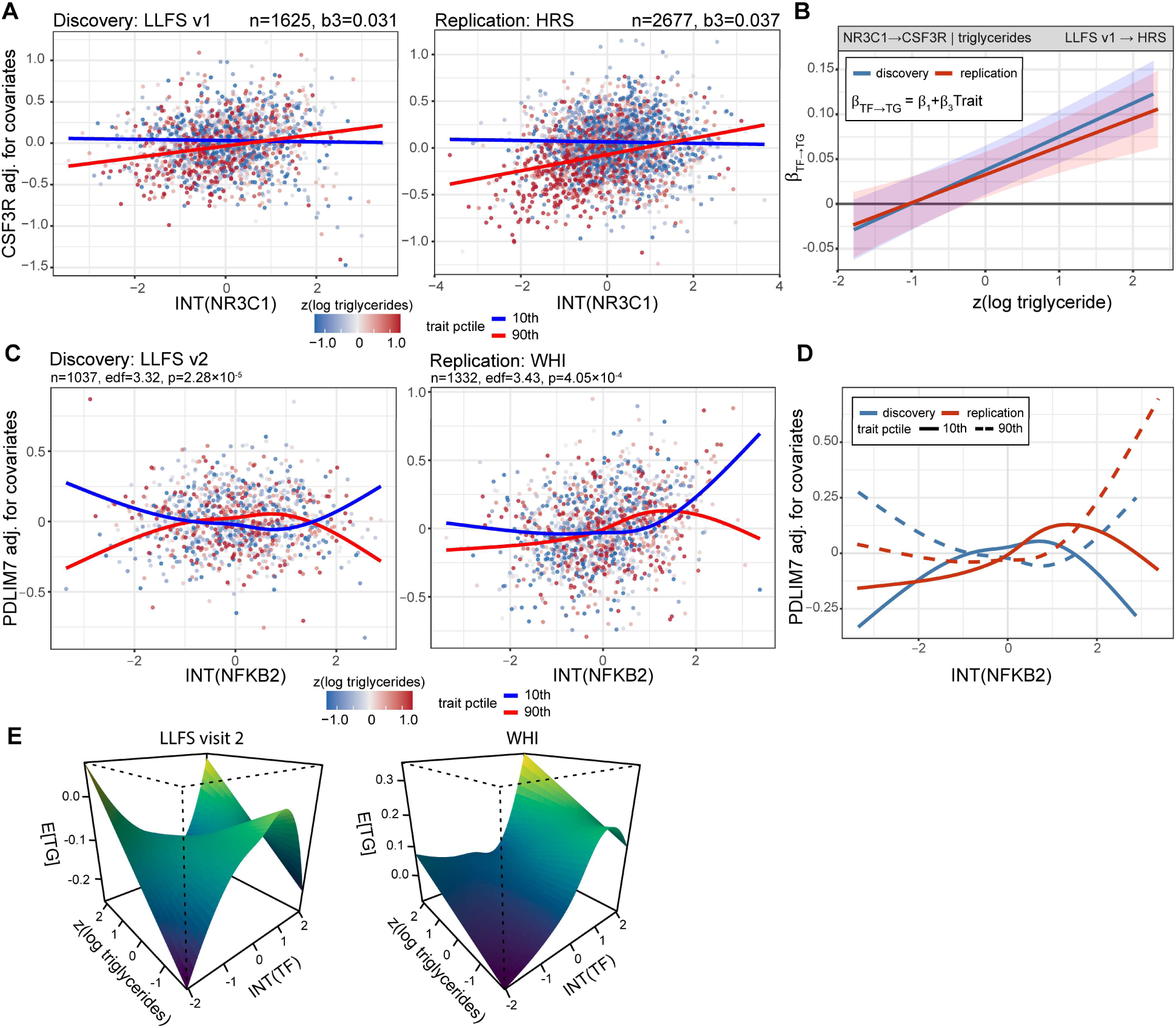
CIDER identifies TF-target regulatory relationships that vary with quantitative health traits. (**A**) CIDER found that NR3C1→CSF3R regulation varies significantly with log-triglyceride level; the effect was discovered in LLFS visit 1 and replicated in HRS. When triglycerides are low (blue), NR3C1 expression has a negligible negative correlation with CSF3R expression; when triglycerides are high (red), the two are clearly positively correlated. (**B**) NR3C1’s regulation of CSF3R as a function of standardized log-triglyceride level. At z(log-triglyceride) = 0, NR3C1 is weakly positively associated with CSF3R; as log-triglyceride rises, this positive coupling strengthens. (**C**) CIDER-GAMM found that NFKB2→PDLIM7 regulation varies significantly with log-triglyceride level; the effect was discovered in LLFS visit 2 and replicated in WHI. When triglyceride levels are low (blue), NFKB2 expression has a concave-down correlation with PDLIM7 expression; when triglycerides are high (red), the two have a concave-up correlation. (**D**) Plots of the fitted NFKB2-PDLIM7 curves at 10 (solid lines) and 90 percentiles (dashed lines) of log-triglycerides for discovery (blue) and replication (red). (**E**) Fitted interaction surfaces for NFKB2-PDLIM7 in LLFS visit 2 and WHI. Expected PDLIM7 expression from the fitted GAM evaluated on a 45 × 45 grid spanning the 2nd-98th percentiles of INT(NFKB2) and z(log triglycerides), with covariates held at their median (continuous) or modal (categorical) values; for LLFS visit 2, the surface is drawn from a plain GAM whereas the interaction statistic quoted in the text is from the kinship GAMM on the same data. The surface is concave in INT(NFKB2) at low triglycerides and convex at high triglycerides.

#### NFKB2→PDLIM7: coupling between the non-canonical NF-κB precursor and a canonical NF-κB brake varies significantly and nonlinearly with circulating triglycerides

CIDER-GAMM identified a regulatory relationship between NFKB2, which encodes the p100/p52 precursor-subunit of the non-canonical NF-κB pathway, and PDLIM7, which encodes a PDZ-LIM adaptor protein that acts as a ubiquitin ligase promoting degradation of the canonical NF-κB subunit p65, whose shape varies significantly and nonlinearly with triglyceride levels (discovery, LLFS: visit 2, *n* = 1037, edf = 3.32, p = 2.3 × 10^-5^; replication, WHI: n = 1332; p = 4.1 × 10^-4^) (Fig. 4C). The interaction is not a change of slope alone: in both cohorts the fitted NFKB2-PDLIM7 relationship is concave downward among individuals with low triglycerides and concave upward among individuals with high triglycerides, a replicated reversal of curvature (Fig. 4D). The linear CIDER independently recovers this edge in three cohorts as a positive, attenuating interaction; the fitted surface shows that this attenuation is the linear projection of a shape change that only the tensor-product smooth function can represent.

## 4 DISCUSSION

We have introduced CIDER, a method that tests whether the regulatory relationship between a transcription factor and its target gene varies with a continuous trait. The central move is to treat the trait as an effect modifier and to detect context-dependent regulation as an interaction in a per-pair regression, rather than as a difference between correlations estimated in two groups. This keeps the trait on its natural, continuous scale, and it extends without difficulty from a linear interaction, which changes the slope of the relationship, to a generalized additive interaction, which changes its shape. In simulations, CIDER outperformed four established two-group methods across sample sizes, effect sizes, and noise levels; applied to whole-blood expression and ten cardiometabolic traits across four independent cohorts, it recovered replicated regulatory relationships that vary with a trait.

The benchmark isolates why the continuous framing helps. By comparing CIDER against a version of itself that sees only the median-dichotomized trait, we found that most of CIDER’s advantage over the two-group methods comes not from the choice of a regression model but simply from declining to discretize the trait. This margin is not fixed: as the trait becomes less separable, and the median split therefore less informative, the advantage of modeling the target’s conditional mean by regression grows in its own right. Because most continuous physiological variables are unimodal rather than cleanly bimodal, this is the regime that matters in practice, and it is precisely where dichotomization is most wasteful.

A separate axis of the benchmark motivates the nonlinear form. When the effect modification changes the shape of the relationship rather than only its slope, some forms cannot be expressed as a difference between the two correlations at all. The clearest case is an interaction in which the transcription factor’s effect inverts from concave to convex across the trait while the within-group correlation stays near zero in both halves: correlation-based methods are blind to it by construction, whereas CIDER-GAM recovers it with a large margin. This is not merely a question of statistical efficiency but of representability; a class of context-dependent regulation that the two-group formulation cannot see motivates having a nonlinear form of the method.

The cohort application shows that these relationships exist in human data and can be replicated across genetically independent cohorts. Reassuringly, the discovered edges include pairs whose direct regulatory link is independently documented, which argues that the recovered edges are real regulatory relationships onto which CIDER adds the trait dependence, rather than statistical artifacts.

The two featured edges also illustrate the kind of mechanistic hypothesis a replicated, trait-dependent edge supports; in keeping with the design of the study, the hypotheses below annotate the statistical findings and played no role in selecting or ranking them.

For NR3C1→CSF3R, if the regulatory shift occurs in response to the lipid environment, one potential mechanistic hypothesis involve transcriptional convergence at the CSF3R promoter. In this model, the baseline inductive effect of NR3C1 is mediated through the CCAAT/enhancer-binding protein (C/EBP) family of transcription factors. Activation of NR3C1 induces the expression and functional engagement of C/EBPα and C/EBPβ [43, 44]. Because the CSF3R promoter region contains canonical C/EBP binding sites, C/EBPα acts as a primary transcriptional activator, establishing basal expression of CSF3R required for myeloid cell survival and granulocytic differentiation [45, 46] By ensuring a steady supply of active C/EBP complexes, NR3C1 keeps the CSF3R promoter engaged, recruiting the necessary chromatin remodeling machinery to hold the DNA in a highly accessible state.

Under the same hypothesis, the amplified regulatory edge observed alongside elevated triglycerides could be driven by pro-inflammatory lipid signaling. High levels of circulating triglycerides and saturated fatty acids are known to initiate intracellular signal transduction cascades analogous to those triggered by potent inflammatory cytokines like TNFα [47, 48]. These cascades culminate in the activation of the NF-κB pathway, which has been shown to drive the nuclear translocation of active subunits such as p65 and RelB in mice, which in turn enhances the mRNA transcription of CSF3R [49].

The interaction detected by CIDER may capture the mechanistic overlap of these two pathways. In a high-triglyceride environment, a lipid-induced surge of NF-κB could enter the nucleus and bind its regulatory motifs efficiently because the CSF3R promoter has already been unmasked and stabilized by the NR3C1-driven C/EBP axis. This simultaneous co-binding of the baseline C/EBP factors and the lipid-driven NF-κB factors would enhancer RNA polymerase II recruitment, amplifying CSF3R transcription. Alternatively, individuals in whom this transcriptional convergence is already highly active may be predisposed to elevated circulating lipids.

For NFKB2→PDLIM7, both genes are components of the NF-κB circuit natively expressed in leukocytes, and their co-expression plausibly reads out the state of a feedback loop. NFKB2 is itself a transcriptional target of canonical NF-κB signaling [50], while PDLIM7 restrains that same canonical branch by promoting turnover of p65 [51]; the pair therefore sits on opposite sides of a negative-feedback relationship between the canonical and non-canonical branches. Lipid exposure engages this circuit directly: free fatty acids and lipid loading activate NF-κB signaling in monocytes and neutrophils, and elevated blood triglycerides are associated with transcriptomic change in whole blood, with genetic evidence consistent with lipids driving a subset of those expression changes rather than merely marking them [52]. A feedback circuit operating at different set points can present qualitatively different input-output geometry, saturating in one regime and accelerating in another, which is the class of behavior a curvature reversal across continuous exposure would produce. We advance this as a hypothesis consistent with the data, not as demonstrated mechanism: no direct experimentally validated NFKB2-PDLIM7 regulatory axis has been reported, and in the one available blood-lineage perturbation, NFKB2 knockdown leaves mean PDLIM7 expression unchanged, consistent with an effect-modification relationship, which a mean-level assay is not designed to detect, rather than with mean regulation. Because NF-κB signaling and lipid metabolism influence each other bidirectionally, the direction of dependence cannot be resolved by an interaction test, and we do not assign one.

An interaction test of this kind is statistically symmetric, and it is important to be explicit about what that does and does not license. Detecting that a TF-target relationship varies with a trait establishes that the two are associated; it does not establish the direction of causation. The regulatory change could contribute to the trait, could reflect it, or both, and an interaction test cannot distinguish these. We therefore describe the discovered edge as regulation that varies with, or depends on, the trait, and we leave open the question of whether a given change contributes causally to the trait – a question that would require longitudinal or interventional data to resolve.

Several limitations bear on how the application results should read. First, whole blood is a mixture of cell types, and a trait associated with a shift in leukocyte composition could in principle produce trait-varying co-expression without any change in regulation within a cell type [53]. Our primary model adjusts for measured blood-cell composition through the complete blood count and for ten expression principal components that absorb unmeasured composition and technical variation. Nonetheless, adjustment at the resolution of a standard differential cannot exclude confounding by cell subsets or activation states it does not resolve, and within-cell-type or single-cell measurement, together with direct functional follow-up of individual edges, remains the appropriate next step for the featured relationships. Second, the discovered edges are conditional on the reference network used to define the candidate set. We regard this less as a confounder than as a boundary of the experiment: because calibration is carried out separately within each (trait, network) combination, the reference network is part of the experimental definition rather than a nuisance variable, and the candidate restriction is a deliberate trade of genome-wide coverage for the multiplicity control that makes interaction testing tractable. Third, the relationship between CIDER edges and a conventional TF perturbation should not be over-read: a knockdown tests whether the TF sets the target’s mean expression, which is a different quantity from effect modification, so a null knockdown is consistent with a genuine CIDER edge rather than evidence against it. Finally, the smooth-term significance values from the additive model are approximate, the cohort data are cross-sectional and observational, and several of the traits are not mutually independent; the linear form of the method, in addition, recovers only the best linear approximation to *β*(*T*) and will misstate the shape of a curved relationship even when it correctly detects one.

Beyond addressing these limitations, the framework invites several extensions. The same interaction test applies unchanged to any continuous context, so tissues and cell types other than whole blood, and continuous variables other than cardiometabolic traits are natural targets. The nonlinear form makes the systematic discovery of non-monotonic context dependence feasible, a class inaccessible to two-group methods. Anchoring individual edges to genetic variation would strengthen causal interpretation. More broadly, CIDER makes context-dependence of regulation a directly testable hypothesis on the natural scale of the context, without first discarding the within-context variation in which graded and reversing modulation lives.

## DATA AND CODE AVAILABILITY

The CIDER software, comprising the linear CIDER, CIDER-GAM, and CIDER-GAMM implementations together with all simulation, benchmark, and analysis code used to produce the results and figures reported here, is openly available at https://doi.org/10.5281/zenodo.22047901. The archive includes test data and instructions for installation, execution, and reproduction of the manuscript’s reported numbers and figures.

The real-transcription-factor resampling benchmarks draw on GTEx v10 whole-blood expression, which is publicly available from the GTEx Portal and is cited in the reference list.

The cohort transcriptome and phenotype data underlying the application (LLFS, FHS, HRS, and WHI) are data from human research participants and are subject to controlled access; the authors are not permitted to redistribute them. These data are available through the database of Genotype and Phenotypes (dbGaP) under the standard Data Access Request and Data Use Agreement process, under study accessions phs000007.v35.p1 (FHS), phs000428.v2.p2 (HRS), and phs001237.v4.p2 (WHI). The analyses reported here were performed under dbGaP project # 33268.

## ACKNOWLEDGEMENTS

We are grateful to the entire Long Life consortium, its participants, and its investigators, without whom this work would not have been possible. The Framingham Heart Study is conducted and supported by the National Heart, Lung, and Blood Institute (NHLBI) in collaboration with Boston University [N01-HC-25195, HHSN268201500001I, 75N92019D00031]. This manuscript was not prepared in collaboration with investigators of the Framingham Heart Study and does not necessarily reflect the opinions or views of the Framingham Heart Study, Boston University, or NHLBI. The HRS (Health and Retirement Study) is sponsored by the National Institute on Aging (grant number NIA U01AG009740) and is conducted by the University of Michigan. The WHI program is funded by the National Heart, Lung, and Blood Institute, National Institutes of Health, U.S. Department of Health and Human Services through contracts HHSN268201100046C, HHSN268201100001C, HHSN268201100002C, HHSN268201100003C, HHSN268201100004C, and HHSN271201100004C. This manuscript was not prepared in collaboration with investigators of the WHI, has not been reviewed and/or approved by the Women’s Health Initiative (WHI), and does not necessarily reflect the opinions of the WHI investigators or the NHLBI.

