## Supplementary Methods for "CIDER: detecting changes in gene regulatory networks that are associated with changes in phenotype"

Michael R. Brent

(314) 362-7238

Campus Box 8510

Washington University

Saint Louis, MO 63130

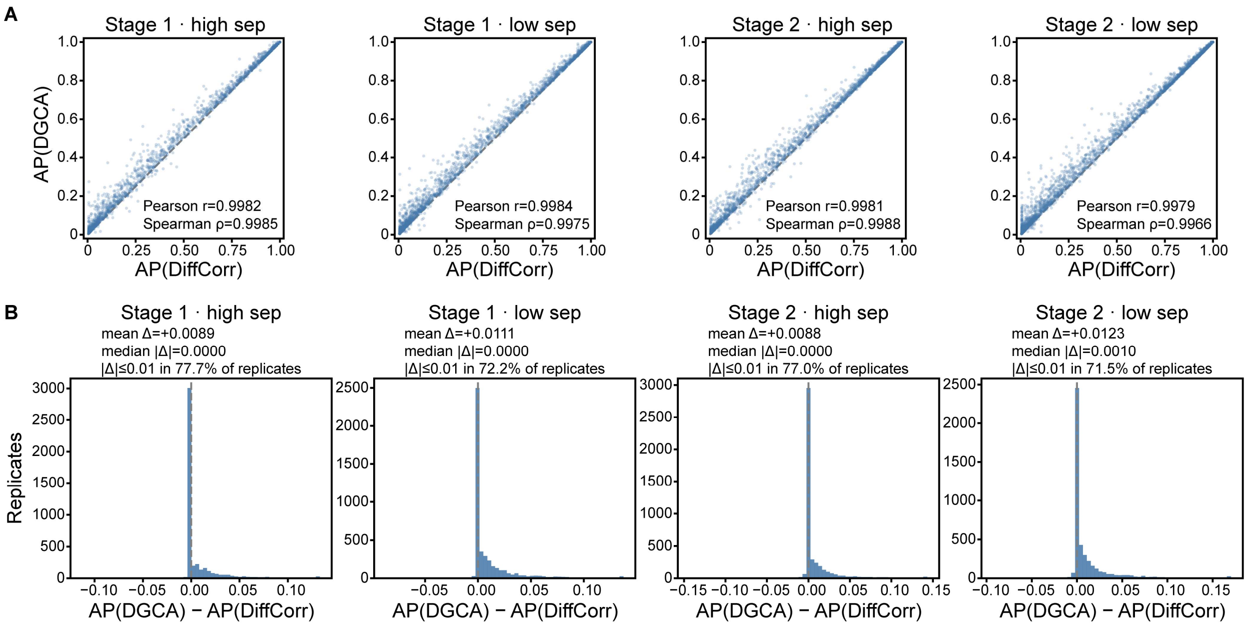


**Supplementary Figure S1. DGCA and DiffCorr performance in stages 1 and 2.** Paired per-replicate average precision (AP) for DGCA and DiffCorr across stages 1 and 2 under high- and low-trait separability. (A) Scatterplots of AP for DGCA and DiffCorr; each point is one simulation replicate (n=4500 per panel). Dashed line is the identity. (B) Distribution of the paired difference $\mathrm{AP(DGCA)} - \mathrm{AP(DiffCorr)}$ over the same replicates; dashed line indicates 0 difference.

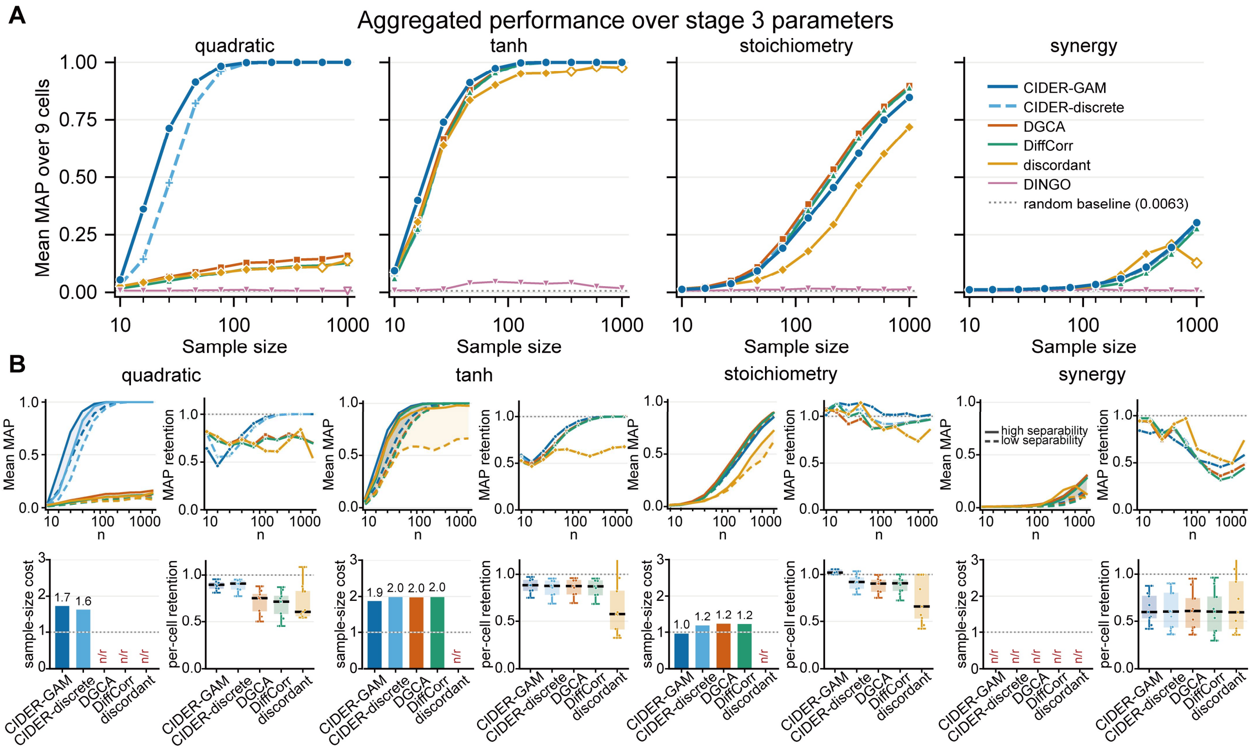


**Supplementary Figure S2. Performance on simulation stage 3.** Performance of CIDER-GAM, CIDER-discrete, DiffCorr, DGCA, and DINGO on simulation stage 3, which has four nonlinear ground-truth interaction forms. (A) Overall performance in stage 3, the mean of the MAP scores across the nine scenarios. (C) For each form, evaluation metrics comparing performance under high versus low trait separability.

**Supplementary Methods**

### S1 Detailed specifications of the alternative two-group methods (main text section 2.10)

All four methods receive the identical dichotomized data: the same TF-target pairs and the same binary group labels obtained by splitting the trait at the sample median.

#### S1.1 DiffCorr and DGCA

DiffCorr and DGCA (Differential Gene Correlation Analysis) both test for a difference in the Pearson correlation of a TF-TG pair between two conditions using the Fisher $z$-transformation [1, 2]. For a pair $(t, g)$, let $r_1$ and $r_2$ denote the within-group Pearson correlations within group 1 (low trait) and group 2 (high trait), with sample sizes $n_1$ and $n_2$. Both methods apply the variance-stabilizing Fisher $z$-transformation,

$$z = \operatorname{atanh}(r) = \frac{1}{2}\log_e\left(\frac{1+r}{1-r}\right),$$

which serves as a normalizing transformation. Under bivariate normality the resulting $z$-score has variance $s^2 = 1/(n_k - 3)$, evaluated separately in each condition to accommodate unequal group sizes and gene-specific missingness, and requires at least four samples per condition. The difference in $z$-scores is standardized by its standard error,

$$Z = \frac{z_1 - z_2}{\sqrt{s_{z_1}^2 + s_{z_2}^2}} = \frac{z_1 - z_2}{\sqrt{\frac{1}{n_1 - 3} + \frac{1}{n_2 - 3}}},$$

and referred to a standard normal distribution under the null hypothesis of equal population correlations, giving a two-sided $p$-value. DiffCorr presents the statistic in the right-hand form and DGCA in the left-hand form (writing it $dz$), but the two are the same test. The methods differ in how pairs are supplied and filtered and in how multiple testing is handled.

#### S1.2 DiffCorr

We computed Pearson correlations for all TF-TG pairs within each group, applied the Fisher $z$-transformation, evaluated $Z$ as above, and obtained two-sided $p$-values from the standard normal distribution. The $p$-values were adjusted by the Benjamini-Hochberg procedure and pairs ranked by the adjusted $p$-value. Because the normal $p$-value is a strictly monotone function of $|Z|$ and the Benjamini-Hochberg adjustment is monotone in the raw $p$-value, this ranking is equivalent to ranking by $|Z|$, and no ties arise.

#### S1.3 DGCA

DGCA takes as input an expression matrix, a design matrix encoding condition membership, and a specification of the two conditions to compare. We built the design matrix from the binarized trait with makeDesign and supplied TF and TG expression as separate input matrices (inputMat and inputMatB), so that DGCA evaluated only TF-TG pairs and not TF-TF or TG-TG pairs, matching the pair set used by every other method. DGCA can optionally filter genes with low expression level or low dispersion before analysis, on the grounds that such genes are more prone to spurious correlations; we did not apply this filter, since the TF-TG pair set was fixed by the simulation design and identical across methods. We ran ddcorAll with corrType = "pearson" and calibrated $p$-values by permuting the condition labels (adjust = "perm", nPerms=100).

Pairs were ranked by the permutation-based differential-correlation $p$-value, with ties broken by the magnitude of the differential $z$-statistic (zScoreDiff). The tiebreak is necessary: with 100 permutations the $p$-value cannot fall below $1/101$, and at moderate to large sample sizes all informative pairs, including every true positive, reach this resolution floor and tie.

#### S1.4 Discordant

Discordant [3] also begins from within-group correlations, but rather than testing each pair independently it models the joint distribution of the Fisher $z$-transformed coefficients $(z_1, z_2)$ across all pairs with a finite mixture model fitted by the EM algorithm. Each marginal $z_k$ is modeled as a mixture of three components corresponding to negative correlation, no correlation, and positive correlation. The joint distribution over the two groups then partitions pairs into a $3 \times 3$ grid of classes; the three diagonal classes are concordant (the pair behaves the same way in both groups) and the six off-diagonal classes are discordant (the pair changes class between groups).

For each pair, Discordant returns the posterior probability of belonging to any discordant class, and pairs were ranked by this probability. We used the three-component parameterization with default EM initialization and convergence settings.

#### S1.5 DINGO

DINGO (differential network analysis in genomics) differs from the preceding methods in estimating *partial* rather than marginal correlations, and so targets conditional dependence given all other genes in the network [4]. DINGO decomposes the group-specific inverse-covariance structure into a global component shared across groups and a group-specific local component: it first fits a global Gaussian graphical model to the pooled data, then models group-specific deviations from it, yielding group-specific partial-correlation matrices $\rho^{(1)}$ and $\rho^{(2)}$. A differential score is computed for each edge,

$$d_{ij} = \frac{\rho_{ij}^{(1)} - \rho_{ij}^{(2)}}{\widehat{se}(\rho_{ij}^{(1)} - \rho_{ij}^{(2)})},$$

where the standard error in the denominator is obtained by bootstrap resampling and edges are ranked by $|d_{ij}|$.

The global-local decomposition lets DINGO distinguish genuine rewiring from differences induced by shared upstream structure, a conceptual advantage over marginal-correlation methods. This advantage comes at a substantial computational cost: the bootstrap must be repeated for every edge, and the underlying graphical-model estimation scales poorly with the number of genes.

### S2 Mathematical formulation of CIDER-GAM smooths (main text section 2.3)

To capture continuous differential regulation of a target gene (TG) by a transcription factor (TF) across varying trait levels, CIDER fits a generalized additive model (GAM). The model decomposes the TG expression response into additive main effects and a pure interaction surface:

$$\mathrm{TG} = \beta_0 + f_1(\mathrm{TF}) + f_2(\mathrm{Trait}) + f_{1,2}(\mathrm{TF}, \mathrm{Trait}) + \epsilon,$$

where $\epsilon = N(0, \sigma^2)$. The functions $f_1, f_2$, and $f_{1,2}$ are constructed using penalized regression splines.

#### S2.1 Thin plate regression splines

The main effects, $f_1(\mathrm{TF})$ and $\mathrm{f}_2(\mathrm{Trait})$, absorb the nonlinear, additive marginal associations. They are constructed using low-rank thin plate regression splines (TPRS) [5].

For a generic univariate predictor $x$ and $n$ observations, a full thin plate spline $f(x)$ minimizes the penalized least squares objective:

$$\min_f \sum_{i=1}^{n} \left(y_i - f(x_i)\right)^2 + \lambda \mathcal{J}_2(f),$$

where $\lambda$ is the smoothing parameter and $\mathcal{J}_2(f)$ is the wiggliness penalty. For a one-dimensional predictor, the penalty is the integrated squares second derivative:

$$\mathcal{J}_2(f) = \int \left(\frac{\partial^2 f}{\partial x^2}\right)^2 dx.$$

The full thin plate spline is parametrized by one radial basis function per data point, which is computationally prohibitive ($O(n^3)$) for large genomic datasets. To resolve this, mgcv constructs a TPRS by forming the full $n \times n$ penalty matrix and applying a Lanczos iteration to perform an eigendecomposition. The basis is truncated to the eigenvectors corresponding to the $k$ largest eigenvalues. We set the basis dimension to $k = 5$. The resulting low-rank approximation preserves the optimal smoothing properties of the full thin plate spline while reducing the computational complexity to $O(k^3)$:

$$f(x) = \delta_j B_j(x),$$

where $B_j(x)$ are the truncated basis functions and $\delta_j$ are the estimated coefficients.

#### S2.2 Tensor product splines

The interaction term, $f_{1,2}(\text{TF}, \text{Trait})$, models how the TF-TG regulatory relationship changes across the trait. Because TF expression and the continuous trait may operate on fundamentally different scales and exhibit different degrees of non-linearity, an isotropic smoothing penalty is inappropriate. CIDER employs a tensor product smooth to construct an anisotropic surface.

Let the marginal basis functions for the TF be $a_i(\text{TF})$ for $i = 1, \dots, I$, and the marginal basis functions for the trait be $b_j(\text{Trait})$ for $j = 1, \dots J$. We define both marginal bases with a dimension of 5 ($I = 5, J = 5$). The tensor product basis is formed by the Kronecker product of these marginal bases:

$$f_{1,2}(\text{TF}, \text{Trait}) = \sum_{i=1}^{I} \sum_{j=1}^{J} \beta_{i,j} a_i(\text{TF}) b_j(\text{Trait}).$$

To estimate the coefficient matrix $\beta$, the model applies separate, dimension-specific penalties to the tensor product space. Let $S_{TF}$ and $S_{Trait}$ be the marginal penalty matrices for the TF and trait, respectively. The full anisotropic penalty matrices are constructed using Kronecker products with identity matrices (**I**):

$$S_{TF}^{*} = S_{TF} \otimes \mathbf{I}_{\text{J}}$$
$$S_{Trait}^{*} = \mathbf{I}_{\text{I}} \otimes S_{Trait}$$

The objective function for the interaction surface incorporates independent smoothing parameters ($\lambda_{TF}, \lambda_{Trait}$):

$$\min_{\beta} \|y - X\beta\|^2 + \lambda_{TF} \beta^T S_{TF}^{*} \beta + \lambda_{Trait} \beta^T S_{Trait}^{*} \beta.$$

These smoothing parameters are optimized via Restricted Maximum Likelihood (REML), allowing the model to apply heavy smoothing to the trait axis while allowing flexibility along the TF axis or vice versa, dictated entirely by the data.

**S2.3 Isolation of the pure interaction**

Standard tensor products (mgcv's te() function) include the main effects of their marginal variables. Because CIDER explicitly includes $f_1(\text{TF})$ and $f_2(\text{Trait})$ as separate terms, using a standard tensor product would result in structural collinearity.

To isolate the differential gene regulation, CIDER uses mgcv's ti() construction. This applies functional analysis of variance (ANOVA) constraints to the tensor product basis. Specifically, the marginal main-effect spaces are projected out of the tensor product space prior to fitting. As a result, $f_{1,2}(\text{TF}, \text{Trait})$ is strictly orthogonal to $f_1$ and $f_2$. Testing the null hypothesis $f_{1,2} = 0$ provides a mathematically rigorous test for the presence of a pure, non-additive interaction between the transcription factor and the trait.
